## Supplemental Figs and Tables for "Sex-biased gene expression is repeatedly masculinized in asexual females"

**Parker et al.**

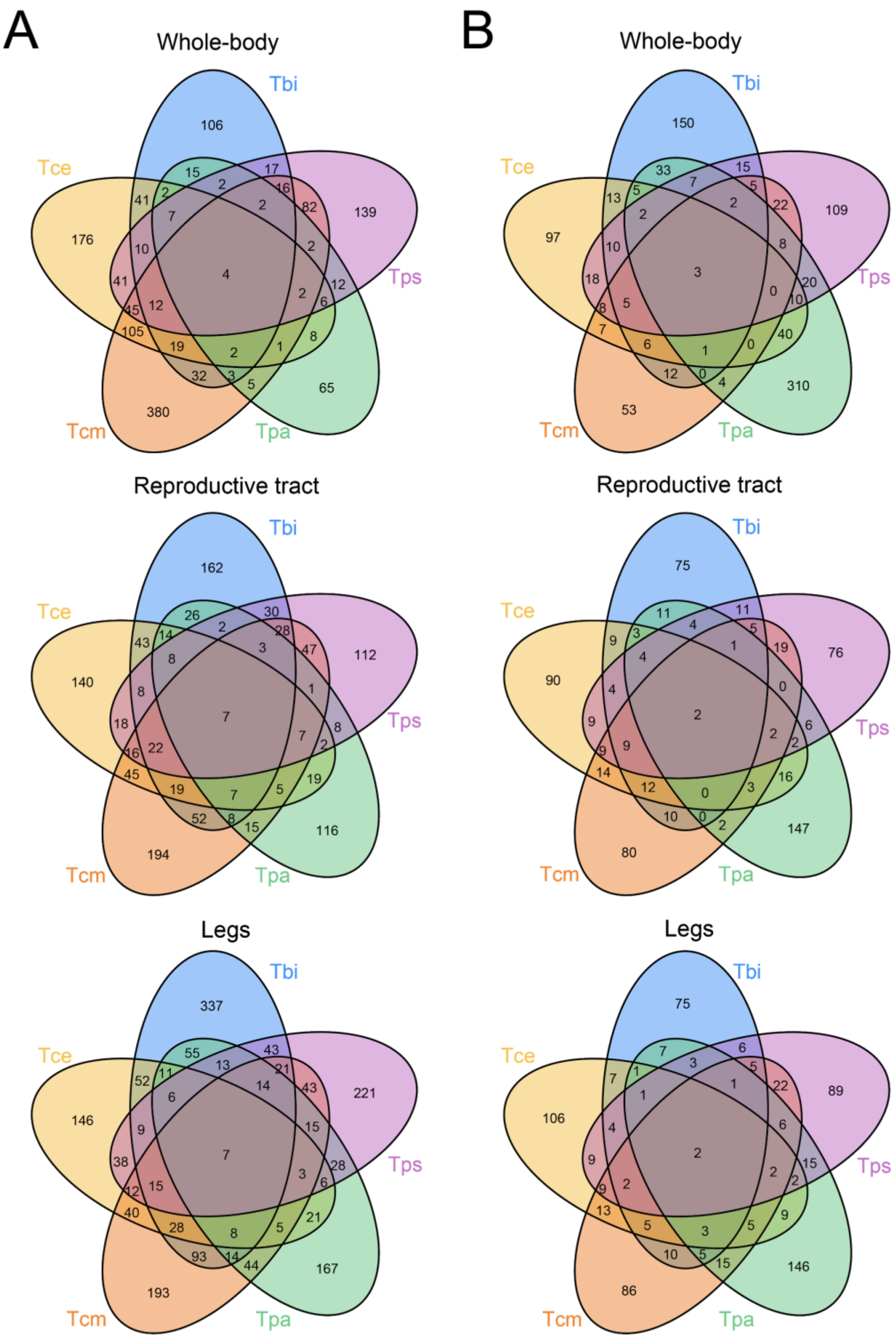


**Supplementary Figure 1 |** Venn-diagrams showing the overlap of enriched GO-terms for (**A**) female-biased and (**B**) male-biased genes. Species names are given as abbreviations: Tbi = *T. bartmani*, Tce = *T.cristinae*, Tps = *T. poppensis*, Tcm = *T. californicum*, Tpa = *T. podura*).

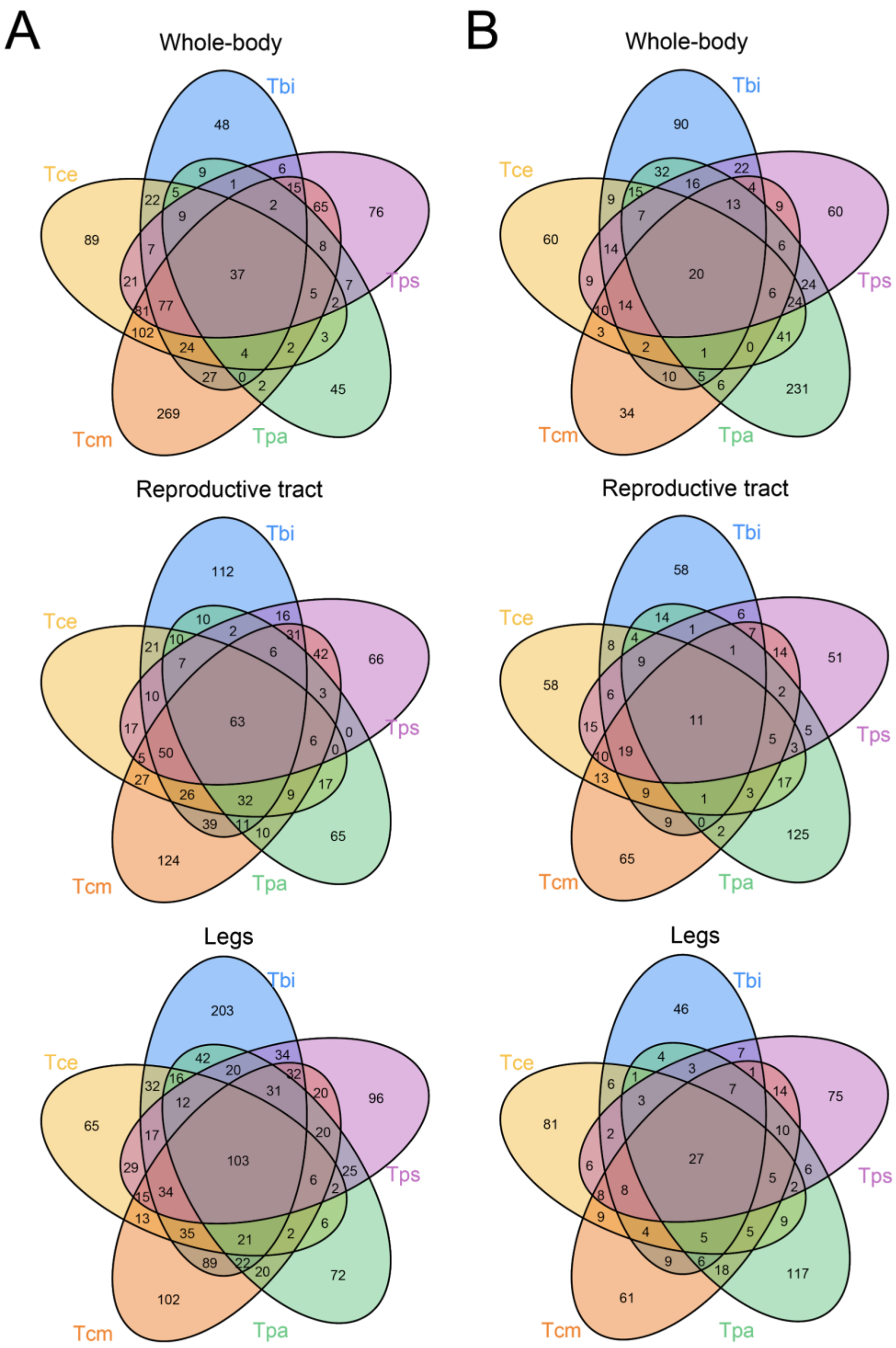


**Supplementary Figure 2** | Venn-diagrams showing the overlap of enriched GO-terms for (**A**) female-biased and (**B**) male-biased genes when GO terms were first clustered together based on parent or child terms. Species names are given as abbreviations: Tbi = *T. bartmani*, Tce = *T. cristinae*, Tps = *T. poppensis*, Tcm = *T. californicum*, Tpa = *T. podura*).

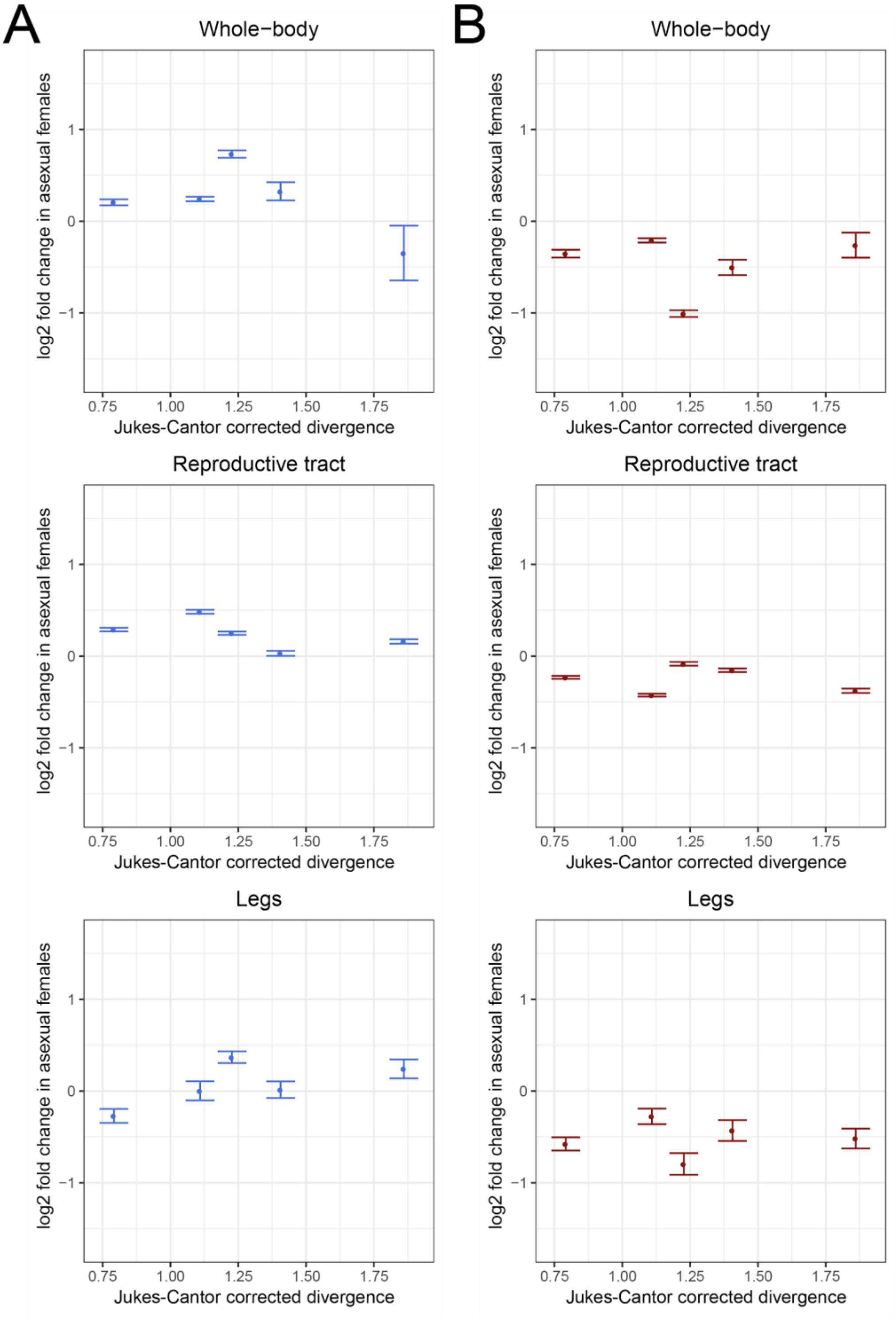


**Supplementary Figure 3** | Relationship between the shift in male-biased (A) or female-biased (B) gene expression in asexual females and sexual-asexual divergence time (estimated as from Jukes–Cantor corrected divergence from Bast et al. [1]) in whole-bodies, reproductive tracts, and legs. Each point is the mean shift in expression for an asexual species with standard error indicated by the error bars.

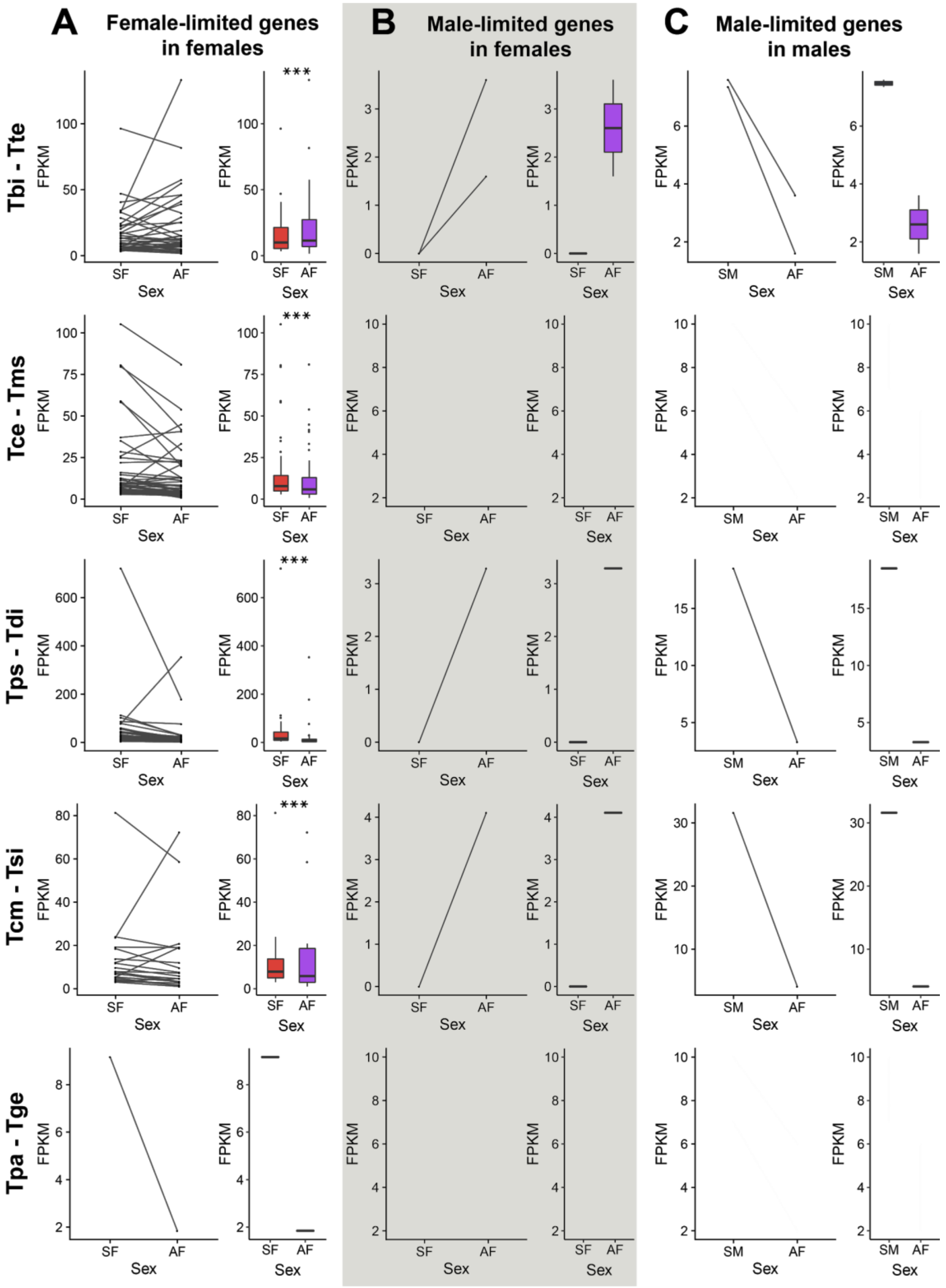


**Supplementary Figure 4 | Expression of sex-limited genes in the whole-body.** A) Expression of female-limited genes in sexual females (SF, red) and asexual females (AF, purple), B) Expression of male-limited genes in sexual females (SF, red) and asexual females

(AF, purple), C) Expression of male-limited genes in sexual males (SM, blue) and asexual females (AF, purple). Asterisks indicate the significance level (FDR) of Wilcoxon tests (*** < 0.001, ** < 0.01, * < 0.05). Species names are given as abbreviations in the form sexual-species - asexual species at the left-hand side (Tbi = *T. bartmani*, Tce = *T.cristinae*, Tps = *T. poppensis*, Tcm = *T. californicum*, Tpa = *T. podura*, Tte = *T. tahoe*, Tms = *T. monikensis*, Tdi = *T. douglasi*, Tsi = *T. shepardi*, and Tge = *T. genevievae*). For the boxplots, boxes represent the interquartile range (25th and 75th percentiles) of the data with the line inside the box representing the median. Whiskers show the most extreme value in the data which is no more than 1.5 times the interquartile range from the box.

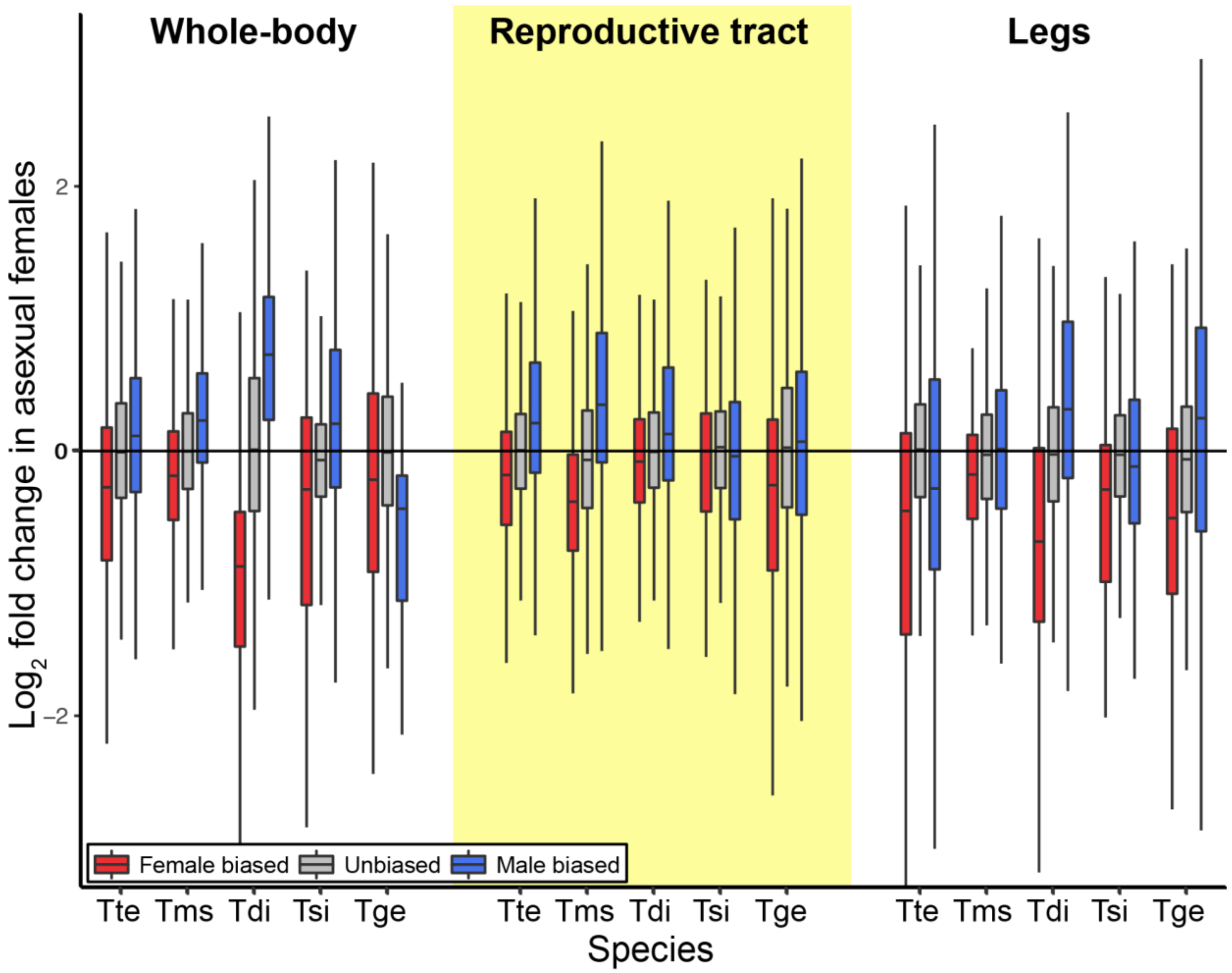


**Supplementary Figure 5 | Expression shift in sex-biased genes in asexual females when including genes with low expression in asexual females.** Positive values on the y-axis indicate increased expression in asexual females. Asterisks indicate the significance level (FDR) of Wilcoxon tests comparing the change in expression in female-biased (red) and male-biased (blue) genes to unbiased genes (*** < 0.001, ** < 0.01, * < 0.05). Species names are abbreviated as follows: Tte = *T. tahoe*, Tms = *T. monikensis*, Tdi = *T. douglasi*, Tsi = *T. shepardi*, and Tge = *T. genevievae*. Boxes represent the interquartile range (25th and 75th percentiles) of the data with the line inside the box representing the median. Whiskers show the most extreme value in the data which is no more than 1.5 times the interquartile range from the box.

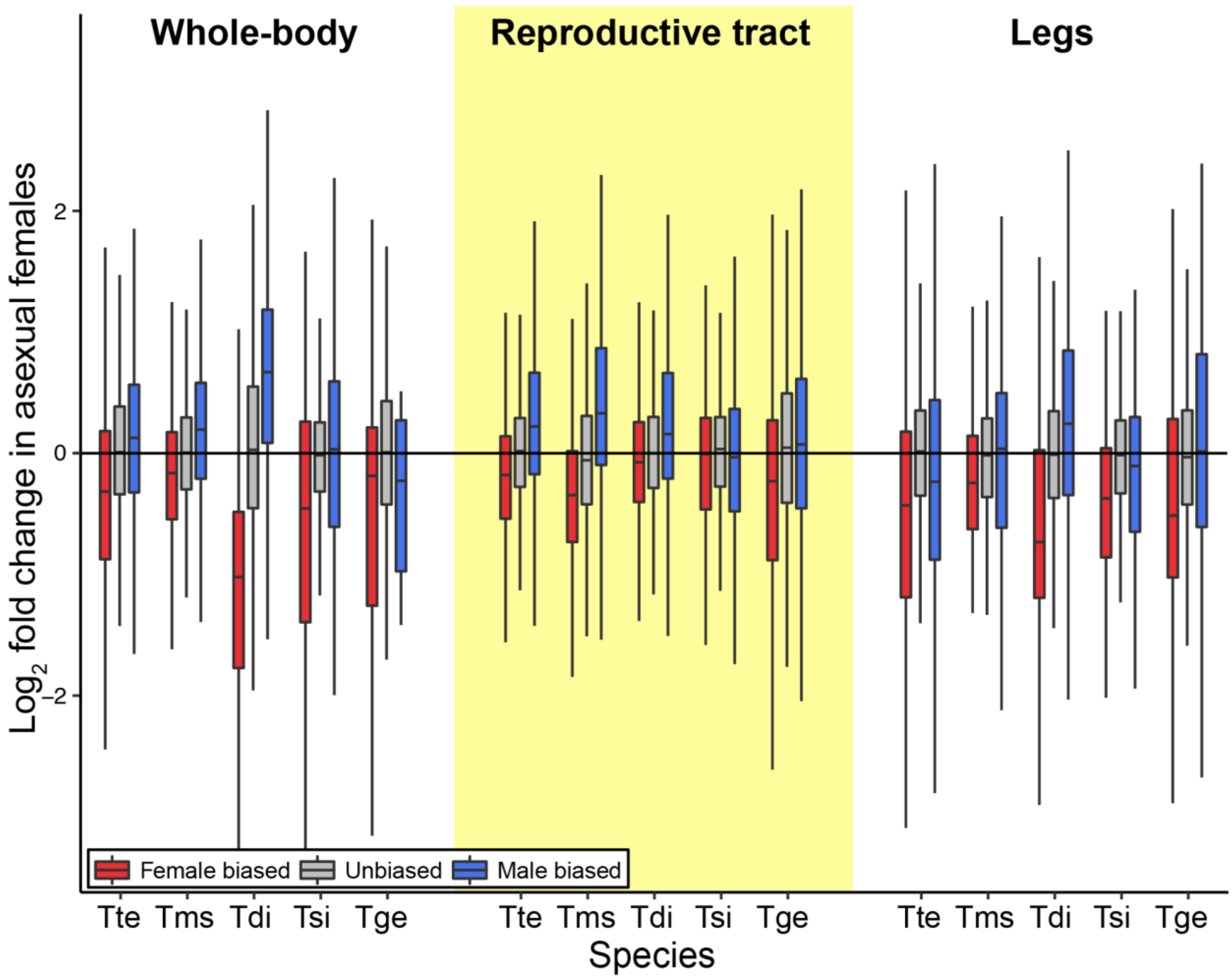


**Supplementary Figure 6 | Expression shift in sex-biased genes in asexual females when using the full sexual-sister species transcriptome as a reference.** Positive values on the y-axis indicate increased expression in asexual females. Species names are abbreviated as follows: Tte = *T. tahoe*, Tms = *T. monikensis*, Tdi = *T. douglasi*, Tsi = *T. shepardi*, and Tge = *T. genevievae*. Boxes represent the interquartile range (25th and 75th percentiles) of the data with the line inside the box representing the median. Whiskers show the most extreme value in the data which is no more than 1.5 times the interquartile range from the box.

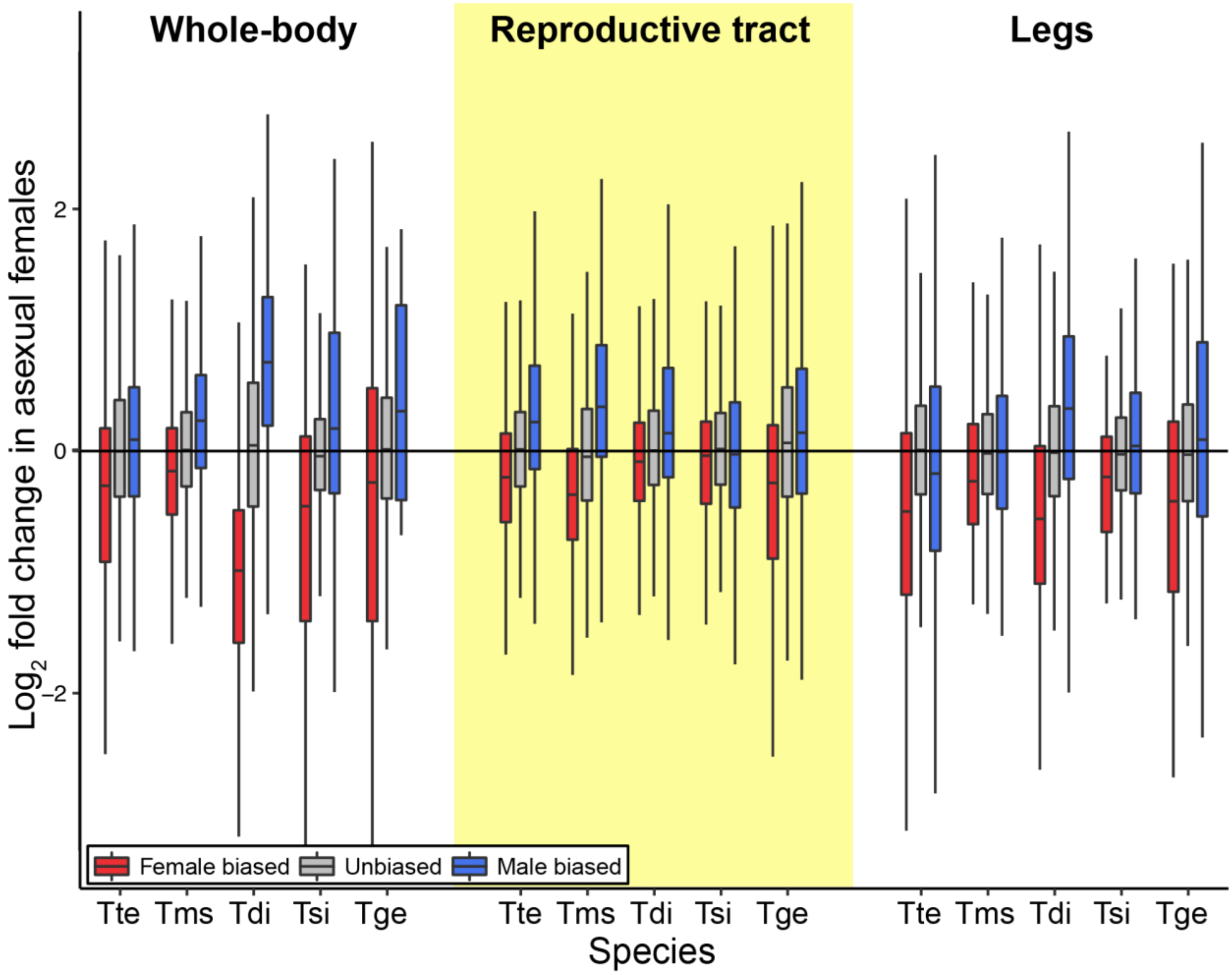


**Supplementary Figure 7 | Expression shift in sex-biased genes in asexual females when using the full asexual-sister species transcriptome as a reference.** Positive values on the y-axis indicate increased expression in asexual females. Species names are abbreviated as follows: Tte = *T. tahoe*, Tms = *T. monikensis*, Tdi = *T. douglasi*, Tsi = *T. shepardi*, and Tge = *T. genevievae*. Boxes represent the interquartile range (25th and 75th percentiles) of the data with the line inside the box representing the median. Whiskers show the most extreme value in the data which is no more than 1.5 times the interquartile range from the box.

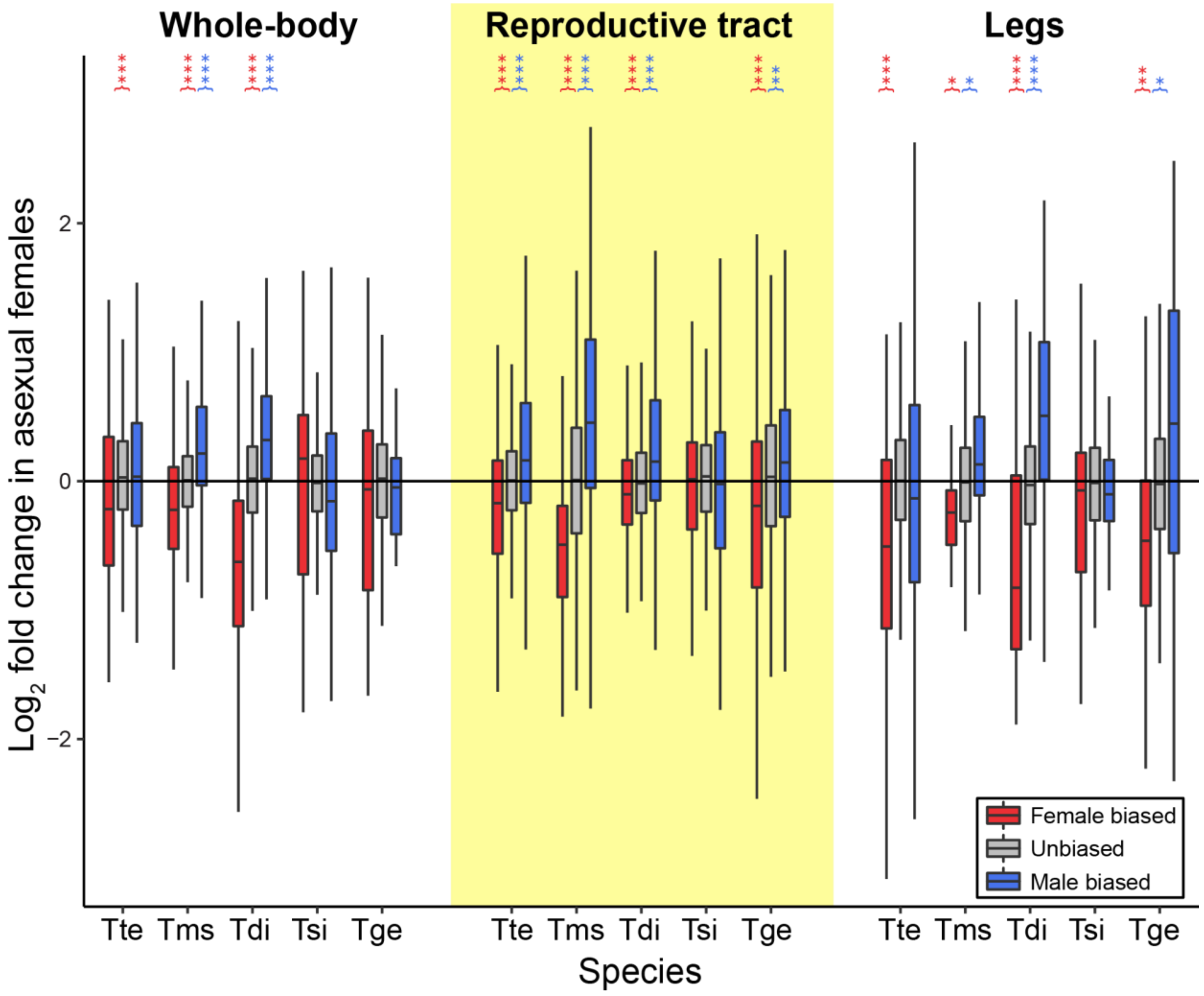


**Supplementary Figure 8 | Expression shift in sex-biased genes in asexual females when using the 10-species orthologs only.** Positive values on the y-axis indicate increased expression in asexual females, negative values indicate decreased expression. Asterisks indicate the significance level (FDR) of Wilcoxon tests comparing the change in expression in female-biased (red) and male-biased (blue) genes to unbiased genes (*** < 0.001, ** < 0.01, * < 0.05). Species names are abbreviated as follows: Tte = *T. tahoe*, Tms = *T. monikensis*, Tdi = *T. douglasi*, Tsi = *T.shepardi*, and Tge = *T. genevievae*. Boxes represent the interquartile range (25th and 75th percentiles) of the data with the line inside the box representing the median. Whiskers show the most extreme value in the data which is no more than 1.5 times the interquartile range from the box.

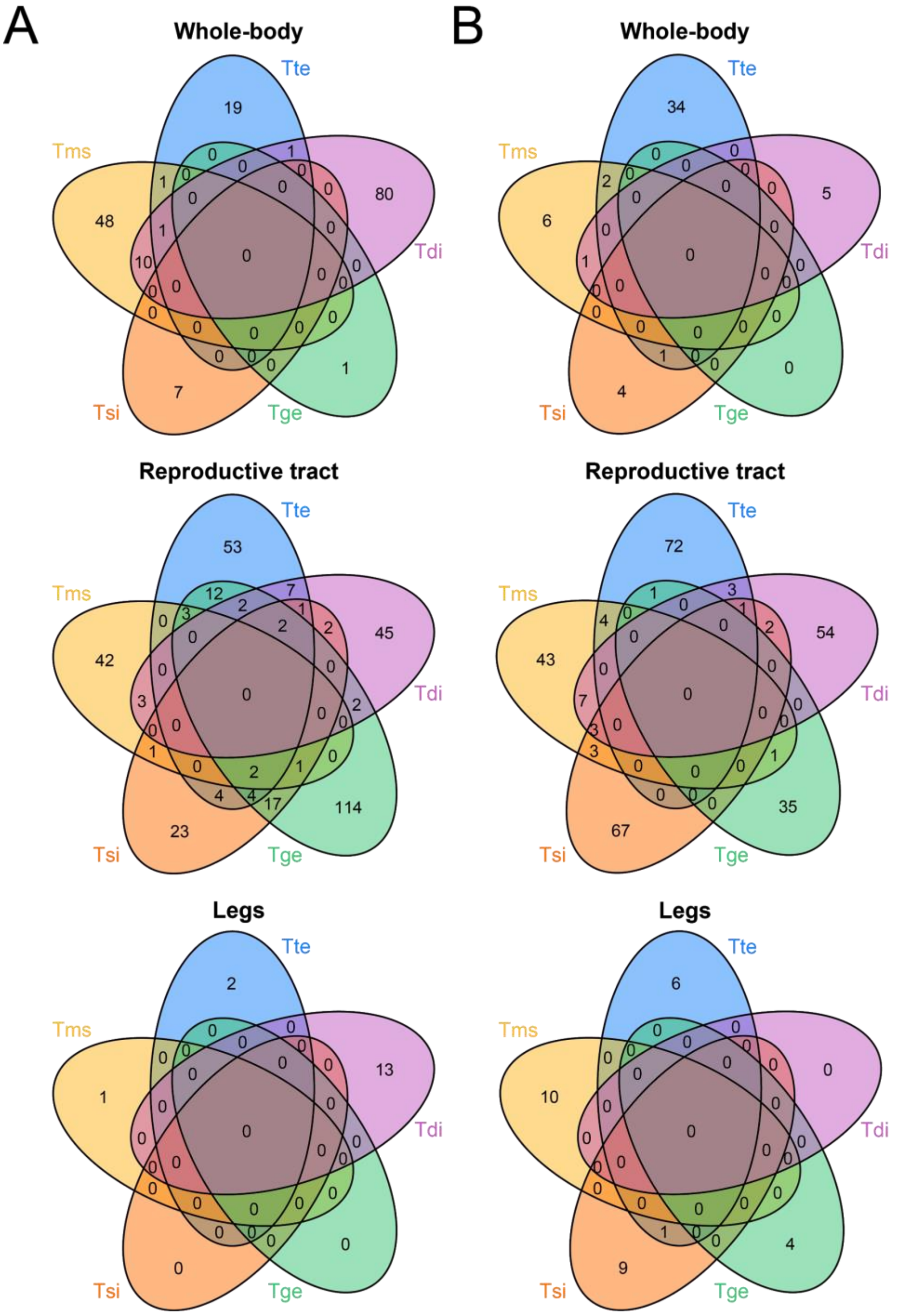


**Supplementary Figure 9 |** Overlap of GO-terms enriched in A) female-biased genes with decreased expression in asexual females, and B) male-biased genes with increased expression in asexual females. Species names are abbreviated as follows: Tte = *T. tahoe*, Tms = *T. monikensis*, Tdi = *T. douglasi*, Tsi = *T.shepardi*, and Tge = *T. genevievae*.

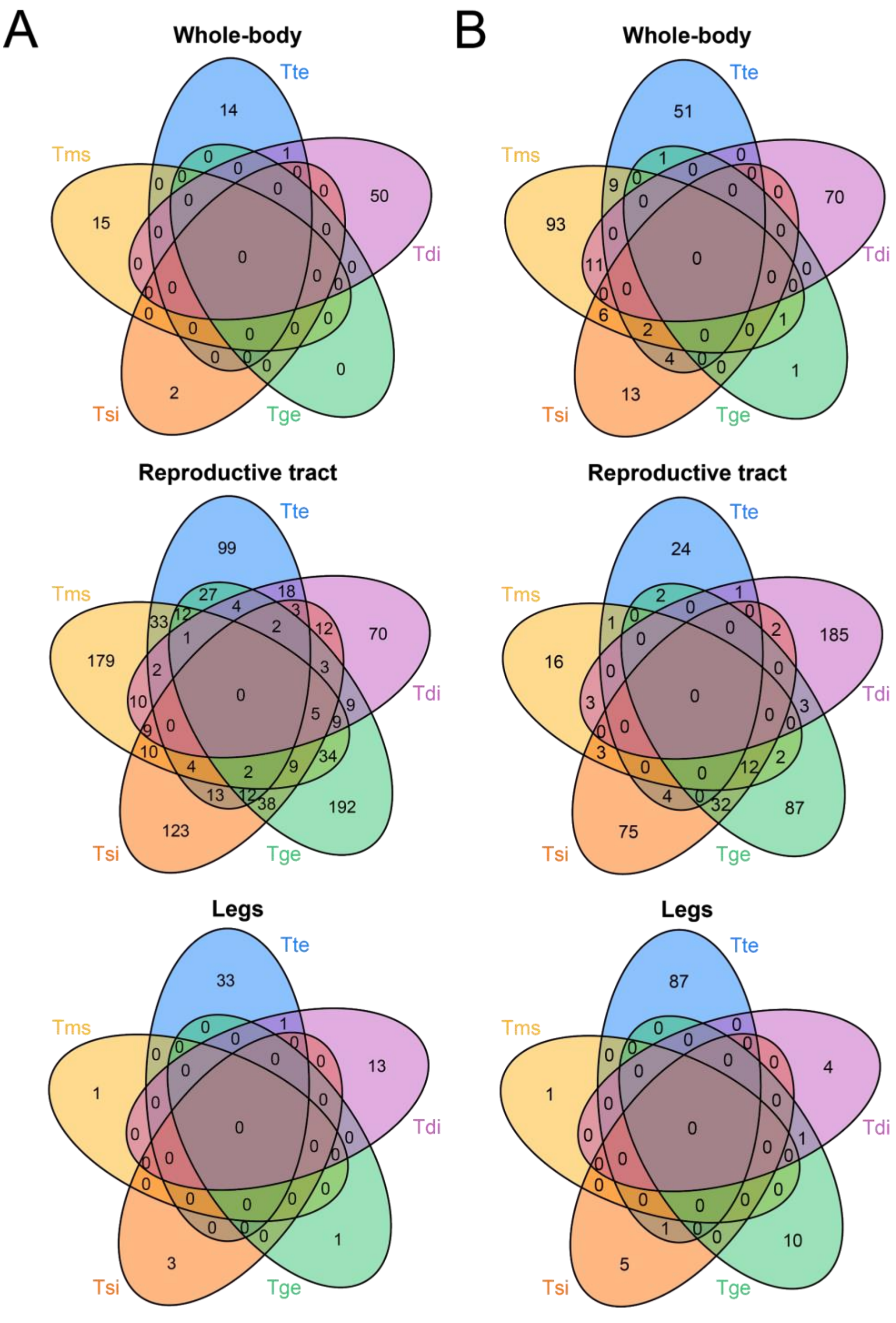


**Supplementary Figure 10 |** Overlap of GO-terms enriched in A) female-biased genes with increased expression in asexual females, and B) male-biased genes with decreased expression in asexual females. Species names are abbreviated as follows: Tte = *T. tahoe*, Tms = *T. monikensis*, Tdi = *T. douglasi*, Tsi = *T.shepardi*, and Tge = *T. genevievae*.

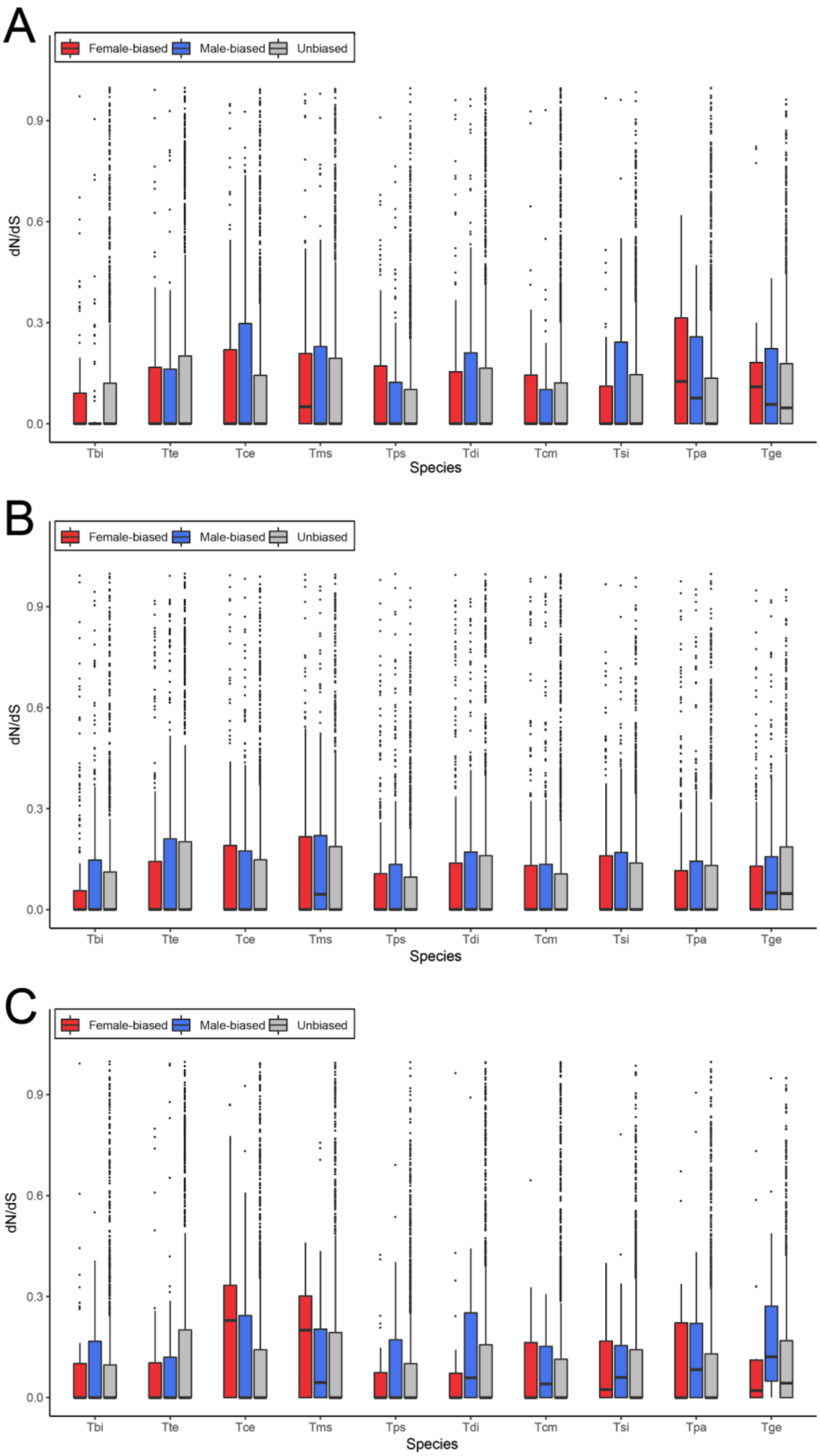


**Supplementary Figure 11 | dN/dS ratios for female-biased (red), male-biased (blue) and unbiased (grey) genes in each of the species.** Species names are abbreviated as follows: Tbi

= *T. bartmani*, Tce = *T. cristinae*, Tps = *T. poppensis*, Tcm = *T. californicum*, Tpa = *T. podura*, Tte = *T. tahoe*, Tms = *T. monikensis*, Tdi = *T. douglasi*, Tsi = *T. shepardi*, and Tge = *T. genevievae.* sexual-asexual sister species pairs are as follows: Tbi-Tte, Tce-Tms, Tps-Tdi, Tcm-Tsi, Tpa-Tge. Boxes represent the interquartile range (25th and 75th percentiles) of the data with the line inside the box representing the median. Whiskers show the most extreme value in the data which is no more than 1.5 times the interquartile range from the box.

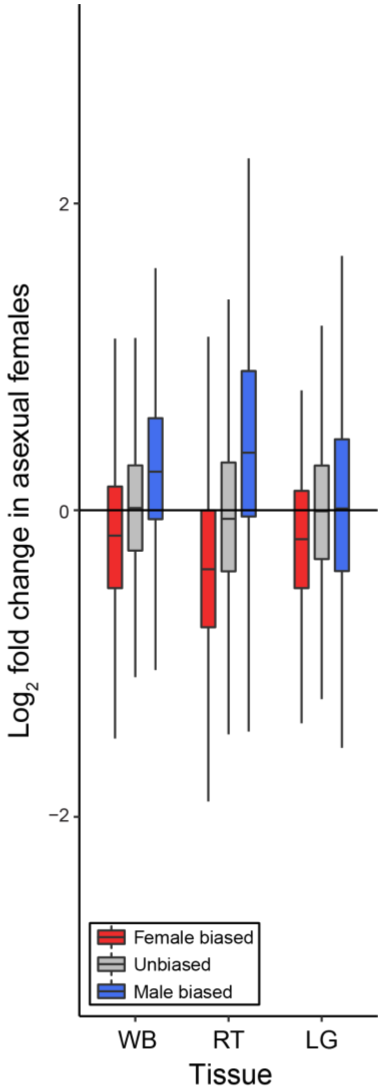


**Supplementary Figure 12 |** Expression shift in autosomal sex-biased genes in asexual female *T. monikensis* in whole-bodies (WB), reproductive tracts (RT) and legs (LG). Boxes represent the interquartile range (25th and 75th percentiles) of the data with the line inside the box representing the median. Whiskers show the most extreme value in the data which is no more than 1.5 times the interquartile range from the box.

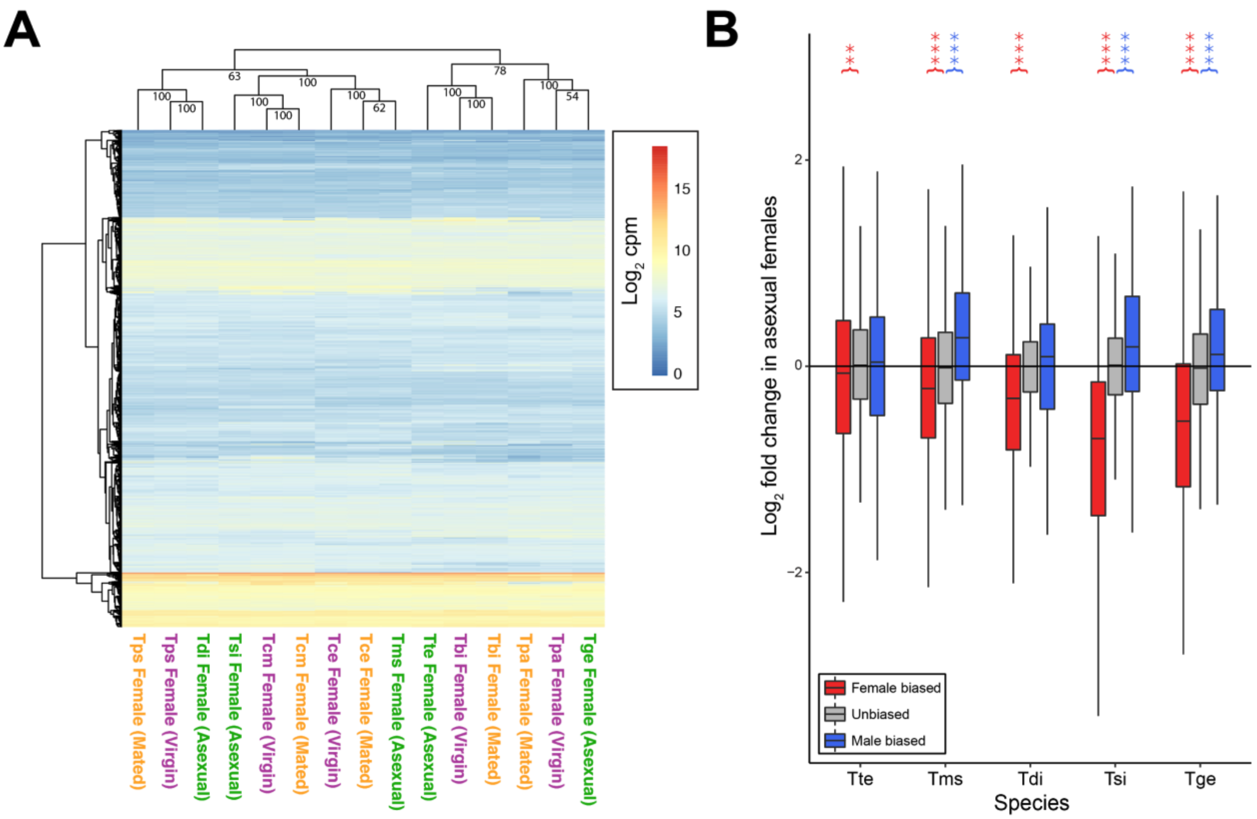


**Supplementary Figure 13** | **A.** Heatmap of expression in whole-bodies shows that whether virgin sexual females have a more similar expression to mated sexual females or asexual females depends upon species. Values on each node show the bootstrap support from 10,000 replicates. **B** Expression shift in sex-biased genes in asexual females in whole-bodies when using virgin sexual female samples. Species names are abbreviated as follows: Tbi = *T. bartmani*, Tce = *T. cristinae*, Tps = *T. poppensis*, Tcm = *T. californicum*, Tpa = *T. podura*, Tte = *T. tahoe*, Tms = *T. monikensis*, Tdi = *T. douglasi*, Tsi = *T. shepardi*, and Tge = *T. genevievae.* Boxes represent the interquartile range (25th and 75th percentiles) of the data with the line inside the box representing the median. Whiskers show the most extreme value in the data which is no more than 1.5 times the interquartile range from the box.

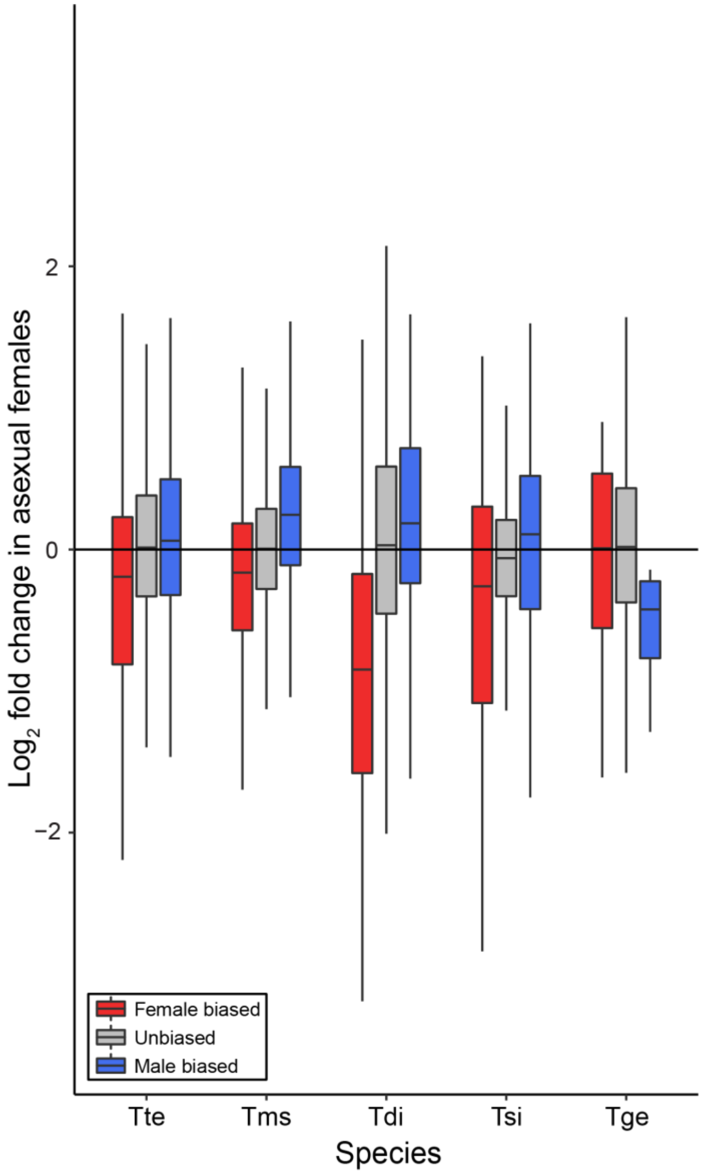


**Supplementary Figure 14 |** Expression shift in sex-biased genes in asexual females in whole-bodies. Sex-biased genes used were identified using both mated and virgin sexual females to independently verify sex-biased genes. Species names are abbreviated as follows: Tbi = *T. bartmani*, Tce = *T. cristinae*, Tps = *T. poppensis*, Tcm = *T. californicum*, Tpa = *T. podura*, Tte = *T. tahoe*, Tms = *T. monikensis*, Tdi = *T. douglasi*, Tsi = *T. shepardi*, and Tge = *T. genevievae.* Boxes represent the interquartile range (25th and 75th percentiles) of the data with the line inside the box representing the median. Whiskers show the most extreme value in the data which is no more than 1.5 times the interquartile range from the box.

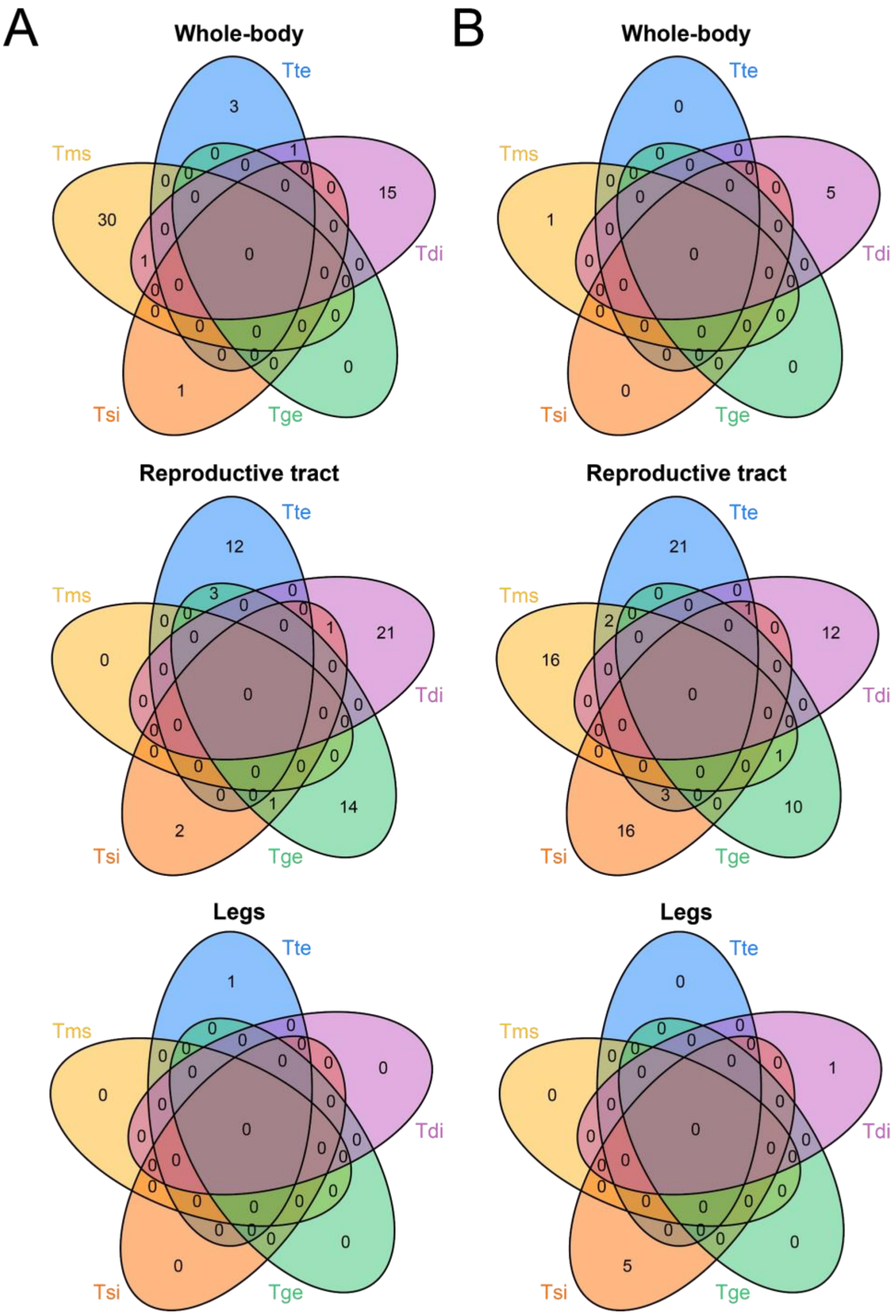


**Supplementary Figure 15 |** Overlap of GO-terms enriched in A) female-biased genes with decreased expression in asexual females, and in B) male-biased genes with increased expression in asexual females using NCBI's nr-arthropod GO annotation. Species names are abbreviated as follows: Tte = *T. tahoe*, Tms = *T. monikensis*, Tdi = *T. douglasi*, Tsi = *T.shepardi*, and Tge = *T. genevievae*.

**Supplementary Table 1 | Numbers of sex-biased genes in each species.** N FB, N MB, and N SB = number of female-, male-, or sex- biased genes respectively. SB dir = sex-bias direction, FDR = adjusted p-value for an excess of male- or female- biased genes. Species names are abbreviated as follows: Tbi = *T. bartmani*, Tce = *T. cristinae*, Tps = *T. poppensis*, Tcm = *T. californicum*, and Tpa = *T. podura*. Tissue types are abbreviated as follows: WB = whole-body, RT = reproductive tract, and LG = legs.

| Species | Tissue | N FB | MB_N | SB_N | SB dir | FDR |
|---|---|---|---|---|---|---|
| **Tbi** | WB | 529 (3.46) | 625 (4.09) | 1154 (7.55) | MB | **0.0083** |
| | RT | 1788 (13.92) | 2057 (16.02) | 3845 (29.94) | MB | **0.0000** |
| | LG | 241 (2) | 342 (2.83) | 583 (4.83) | MB | **0.0001** |
| **Tce** | WB | 946 (6.37) | 846 (5.7) | 1792 (12.07) | FB | **0.0234** |
| | RT | 1863 (15.22) | 1734 (14.16) | 3597 (29.38) | FB | **0.0360** |
| | LG | 69 (0.63) | 117 (1.07) | 186 (1.7) | MB | **0.0010** |
| **Tps** | WB | 697 (5.03) | 479 (3.46) | 1176 (8.49) | FB | **0.0000** |
| | RT | 1656 (14.1) | 2020 (17.2) | 3676 (31.29) | MB | **0.0000** |
| | LG | 131 (1.17) | 219 (1.95) | 350 (3.12) | MB | **0.0000** |
| **Tcm** | WB | 168 (1.2) | 121 (0.87) | 289 (2.07) | FB | **0.0088** |
| | RT | 1346 (11.82) | 1497 (13.14) | 2843 (24.96) | MB | **0.0083** |
| | LG | 50 (0.45) | 115 (1.03) | 165 (1.48) | MB | **0.0000** |
| **Tpa** | WB | 62 (0.45) | 5 (0.04) | 67 (0.48) | FB | **0.0000** |
| | RT | 1909 (17.24) | 1755 (15.85) | 3664 (33.1) | FB | **0.0146** |
| | LG | 70 (0.71) | 108 (1.09) | 178 (1.8) | MB | **0.0083** |

**Supplementary Table 2 | Common biological coefficient of variation across male and female samples for each sexual species.**

| Species | Whole-body | Rep. tract | Legs |
|---|---|---|---|
| *T. bartmani* | 0.358 | 0.186 | 0.153 |
| *T. cristinae* | 0.324 | 0.415 | 0.294 |
| *T. poppensis* | 0.418 | 0.241 | 0.278 |
| *T. californicum* | 0.483 | 0.375 | 0.349 |
| *T. podura* | 0.426 | 0.317 | 0.290 |

**Supplementary Table 3 |** Wilcoxon tests for expression shifts in sex-biased genes in asexual females. ΔFB and ΔMB are the median shifts of female-biased and male-biased expression in asexual females respectively. Tissue types are abbreviated as follows: WB = whole-body, RT = reproductive tract, and LG = legs. Species names are abbreviated as follows: Tte = *T. tahoe*, Tms = *T. monikensis*, Tdi = *T. douglasi*, Tsi = *T. shepardi*; Tge = *T. genevievae.*

| Sp | Tissue | ΔFB | FB FDR | ΔMB | MB FDR |
|---|---|---|---|---|---|
| Tte | WB | -0.249 | **1.11 x $10^{-25}$** | 0.139 | **9.72 x $10^{-06}$** |
| Tms | WB | -0.178 | **1.24 x $10^{-27}$** | 0.245 | **9.83 x $10^{-39}$** |
| Tdi | WB | -0.848 | **3.50 x $10^{-193}$** | 0.749 | **2.48 x $10^{-65}$** |
| Tsi | WB | -0.283 | **9.00 x $10^{-06}$** | 0.223 | **3.48 x $10^{-06}$** |
| Tge | WB | -0.184 | 0.0755 | -0.252 | 0.2749 |
| Tte | RT | -0.179 | **3.50 x $10^{-50}$** | 0.223 | **1.30 x $10^{-70}$** |
| Tms | RT | -0.377 | **1.68 x $10^{-110}$** | 0.379 | **8.08 x $10^{-142}$** |
| Tdi | RT | -0.071 | **3.00 x $10^{-09}$** | 0.147 | **4.40 x $10^{-33}$** |
| Tsi | RT | -0.009 | **2.14 x $10^{-06}$** | -0.025 | **0.0090** |
| Tge | RT | -0.252 | **1.01 x $10^{-61}$** | 0.111 | **0.0016** |
| Tte | LG | -0.421 | **1.30 x $10^{-18}$** | -0.232 | **0.0016** |
| Tms | LG | -0.169 | **0.0045** | 0.051 | 0.3360 |
| Tdi | LG | -0.654 | **1.63 x $10^{-14}$** | 0.314 | **3.84 x $10^{-12}$** |
| Tsi | LG | -0.291 | **6.24 x $10^{-05}$** | -0.054 | 0.8031 |
| Tge | LG | -0.503 | **2.14 x $10^{-06}$** | 0.277 | **0.0014** |

**Supplementary Table 4 |** Wilcoxon tests for changes in expression in female-limited genes. SM = expression (FPKM) in males, SF = expression (FPKM) in sexual females, AF = expression (FPKM) in asexual females. N genes = Number of female-limited genes. Tissue types are abbreviated as follows: WB = whole-body, RT = reproductive tract, and LG = legs. Species names are abbreviated by the sexual species name as follows (sexual species – asexual species): Tbi = *T. bartmani - T. tahoe*; Tce = *T. cristinae - T. monikensis*; Tps = *T. poppensis - T. douglasi*; Tcm = *T. californicum - T. shepardi*; Tpa = *T. podura - T. genevievae.*

| **Species** | **Tissue** | **SM exp** | **SF exp** | **AF exp** | **N genes** | **FDR** |
|---|---|---|---|---|---|---|
| Tbi | WB | 0 | 9.93 | 11.33 | 39 | **7.64 x 10-11** |
| Tce | WB | 0 | 7.83 | 5.87 | 50 | **4.09 x 10-09** |
| Tps | WB | 0 | 16.51 | 6.44 | 50 | **4.09 x 10-09** |
| Tcm | WB | 0 | 7.86 | 5.83 | 21 | **2.86 x 10-06** |
| Tpa | WB | 0 | 9.16 | 1.85 | 1 | 1 |
| Tbi | RT | 0 | 13.85 | 10.30 | 33 | **2.44 x 10-09** |
| Tce | RT | 0 | 20.06 | 9.07 | 29 | **1.05 x 10-05** |
| Tps | RT | 0 | 11.43 | 4.46 | 13 | **4.66 x 10-04** |
| Tcm | RT | 0 | 21.10 | 6.38 | 15 | **1.28 x 10-04** |
| Tpa | RT | 0 | 44.34 | 19.62 | 30 | **7.82 x 10-09** |
| Tbi | LG | 0 | 4.75 | 0.020 | 1 | 1 |
| Tce | LG | NA | NA | NA | 0 | NA |
| Tps | LG | NA | NA | NA | 0 | NA |
| Tcm | LG | NA | NA | NA | 0 | NA |
| Tpa | LG | NA | NA | NA | 0 | NA |

**Supplementary Table 5 |** Wilcoxon tests for changes in expression in male-limited genes. SM = expression (FPKM) in males, SF = expression (FPKM) in sexual females, AF = expression (FPKM) in asexual females. N genes = Number of male-limited genes. Tissue types are abbreviated as follows: WB = whole-body, RT = reproductive tract, and LG = legs. Species names are abbreviated by the sexual species name as follows (sexual species – asexual species): Tbi = *T. bartmani - T. tahoe*; Tce = *T. cristinae - T. monikensis*; Tps = *T. poppensis - T. douglasi*; Tcm = *T. californicum - T. shepardi*; Tpa = *T. podura - T. genevievae.*

| Species | Tissue | SM exp | SF exp | AF exp | N genes | FDR |
|---|---|---|---|---|---|---|
| Tbi | WB | 7.48 | 0 | 2.60 | 2 | 0.70 |
| Tce | WB | NA | NA | NA | 0 | NA |
| Tps | WB | 18.51 | 0 | 3.29 | 1 | 1 |
| Tcm | WB | 31.59 | 0 | 4.11 | 1 | 1 |
| Tpa | WB | NA | NA | NA | 0 | NA |
| Tbi | RT | 9.85 | 0 | 0.07 | 26 | **$1.04 \times 10^{-07}$** |
| Tce | RT | 7.14 | 0 | 0.12 | 17 | **$3.56 \times 10^{-05}$** |
| Tps | RT | 7.69 | 0 | 0.04 | 11 | **$1.58 \times 10^{-03}$** |
| Tcm | RT | 14.45 | 0 | 0 | 5 | 0.47 |
| Tpa | RT | 11.03 | 0 | 0.25 | 11 | **$1.58 \times 10^{-03}$** |
| Tbi | LG | 7.26 | 0 | 0 | 1 | 1 |
| Tce | LG | NA | NA | NA | 0 | NA |
| Tps | LG | NA | NA | NA | 0 | NA |
| Tcm | LG | 4.52 | 0 | 0 | 1 | 1 |
| Tpa | LG | NA | NA | NA | 0 | NA |

**Supplementary Table 6 | Number of male-biased genes excluded due to low expression in asexual females.** MB_excl = number of male-biased genes excluded, total_excl = total number of genes excluded, MB = number of male-biased genes in the main analysis, total = number of genes in the main analysis, fisher_OR = odd-ratio from a Fisher's exact test, fisher_FDR = FDR value from the Fisher's exact test. Tissue types are abbreviated as follows: WB = whole-body, RT = reproductive tract, and LG = legs. Species names are abbreviated by the sexual species name as follows (sexual species – asexual species): Tbi = *T. bartmani - T. tahoe*; Tce = *T. cristinae - T. monikensis*; Tps = *T. poppensis - T. douglasi*; Tcm = *T. californicum - T. shepardi*; Tpa = *T. podura - T. genevievae.*

| Sp | Tiss | MB_excl | total_excl | MB | total | fisher_OR | fisher_FDR |
|---|---|---|---|---|---|---|---|
| Tbi | WB | 7 | 119 | 625 | 15282 | 1.47 | 0.5180 |
| Tce | WB | 14 | 240 | 846 | 14842 | 1.02 | 0.9517 |
| Tcm | WB | 2 | 195 | 121 | 13959 | 1.19 | 0.7935 |
| Tpa | WB | 1 | 200 | 5 | 13911 | 13.97 | 0.1368 |
| Tps | WB | 3 | 144 | 479 | 13850 | 0.59 | 0.6161 |
| Tbi | RT | 67 | 225 | 2057 | 12843 | 2.22 | **0.0000** |
| Tce | RT | 59 | 241 | 1734 | 12243 | 1.96 | **0.0001** |
| Tcm | RT | 55 | 266 | 1497 | 11390 | 1.72 | **0.0017** |
| Tpa | RT | 97 | 334 | 1755 | 11070 | 2.17 | **0.0000** |
| Tps | RT | 59 | 230 | 2020 | 11747 | 1.66 | **0.0031** |
| Tbi | LG | 25 | 258 | 342 | 12077 | 3.68 | **0.0000** |
| Tce | LG | 3 | 172 | 117 | 10948 | 1.64 | 0.5951 |
| Tcm | LG | 9 | 206 | 115 | 11143 | 4.38 | **0.0013** |
| Tpa | LG | 7 | 651 | 108 | 9914 | 0.99 | 1.0000 |
| Tps | LG | 10 | 269 | 219 | 11214 | 1.94 | 0.1327 |

**Supplementary Table 7 | Number of female-biased genes excluded due to low expression in asexual females.** FB_excl = number of female-biased genes excluded, total_excl = total number of genes excluded, FB = number of female-biased genes in the main analysis, total = number of genes in the main analysis, fisher_OR = odd-ratio from a Fisher's exact test, fisher_FDR = FDR value from the Fisher's exact test. Tissue types are abbreviated as follows: WB = whole-body, RT = reproductive tract, and LG = legs. Species names are abbreviated by the sexual species name as follows (sexual species – asexual species): Tbi = *T. bartmani - T. tahoe*; Tce = *T. cristinae - T. monikensis*; Tps = *T. poppensis - T. douglasi*; Tcm = *T. californicum - T. shepardi*; Tpa = *T. podura - T. genevievae.*

| Sp | Tiss | FB_excl | total_excl | FB | total | fisher_OR | fisher_FDR |
|---|---|---|---|---|---|---|---|
| Tbi | WB | 1 | 119 | 529 | 15282 | 0.24 | 0.3279 |
| Tce | WB | 2 | 240 | 946 | 14842 | 0.12 | **0.0002** |
| Tcm | WB | 1 | 195 | 168 | 13959 | 0.42 | 0.9163 |
| Tpa | WB | 1 | 200 | 62 | 13911 | 1.12 | 0.8099 |
| Tps | WB | 1 | 144 | 697 | 13850 | 0.13 | **0.0269** |
| Tbi | RT | 12 | 225 | 1788 | 12843 | 0.35 | **0.0002** |
| Tce | RT | 7 | 241 | 1863 | 12243 | 0.17 | **0.0000** |
| Tcm | RT | 6 | 266 | 1346 | 11390 | 0.17 | **0.0000** |
| Tpa | RT | 13 | 334 | 1909 | 11070 | 0.19 | **0.0000** |
| Tps | RT | 27 | 230 | 1656 | 11747 | 0.81 | 0.5083 |
| Tbi | LG | 9 | 258 | 241 | 12077 | 1.78 | 0.2374 |
| Tce | LG | 1 | 172 | 69 | 10948 | 0.92 | 1.0000 |
| Tcm | LG | 0 | 206 | 50 | 11143 | 0.00 | 1.0000 |
| Tpa | LG | 1 | 651 | 70 | 9914 | 0.22 | 0.2455 |
| Tps | LG | 3 | 269 | 131 | 11214 | 0.95 | 1.0000 |

**Supplementary Table 8 | Number of male-biased genes excluded due to low expression in asexual females when using the sexual reference.** MB_excl = number of male-biased genes excluded, total_excl = total number of genes excluded, MB = number of male-biased genes in the main analysis, total = number of genes in the main analysis, fisher_OR = odd-ratio from a Fisher's exact test, fisher_FDR = FDR value from the Fisher's exact test. Tissue types are abbreviated as follows: WB = whole-body, RT = reproductive tract, and LG = legs. Species names are abbreviated by the sexual species name as follows (sexual species – asexual species): Tbi = *T. bartmani - T. tahoe*; Tce = *T. cristinae - T. monikensis*; Tps = *T. poppensis - T. douglasi*; Tcm = *T. californicum - T. shepardi*; Tpa = *T. podura - T. genevievae.*

| sp | Tiss | MB_excl | total_excl | MB_kept | total_kept | MB_fisher_OR | MB_fisher_FDR |
|---|---|---|---|---|---|---|---|
| Tbi | WB | 93 | 758 | 969 | 19426 | 2.66 | **0.0000** |
| Tce | WB | 110 | 1069 | 1116 | 19757 | 1.92 | **0.0000** |
| Tcm | WB | 21 | 704 | 160 | 17661 | 3.36 | **0.0000** |
| Tpa | WB | 1 | 1383 | 6 | 22296 | 2.69 | 0.3684 |
| Tps | WB | 31 | 872 | 560 | 17862 | 1.14 | 0.4865 |
| Tbi | GN | 190 | 602 | 2634 | 16105 | 2.36 | **0.0000** |
| Tce | GN | 118 | 534 | 2292 | 16019 | 1.70 | **0.0000** |
| Tcm | GN | 108 | 500 | 1843 | 14412 | 1.88 | **0.0000** |
| Tpa | GN | 215 | 810 | 2644 | 16838 | 1.94 | **0.0000** |
| Tps | GN | 137 | 583 | 2584 | 15242 | 1.50 | **0.0001** |
| Tbi | LG | 75 | 654 | 474 | 15161 | 4.01 | **0.0000** |
| Tce | LG | 9 | 512 | 171 | 14761 | 1.53 | 0.2401 |
| Tcm | LG | 19 | 584 | 168 | 13733 | 2.71 | **0.0003** |
| Tpa | LG | 28 | 1602 | 164 | 15022 | 1.61 | **0.0325** |
| Tps | LG | 30 | 683 | 324 | 14497 | 2.01 | **0.0013** |

**Supplementary Table 9 | Number of female-biased genes excluded due to low expression in asexual females when using the sexual reference.** FB_excl = number of female-biased genes excluded, total_excl = total number of genes excluded, FB = number of female-biased genes in the main analysis, total = number of genes in the main analysis, fisher_OR = odd-ratio from a Fisher's exact test, fisher_FDR = FDR value from the Fisher's exact test. Tissue types are abbreviated as follows: WB = whole-body, RT = reproductive tract, and LG = legs. Species names are abbreviated by the sexual species name as follows (sexual species – asexual species): Tbi = *T. bartmani - T. tahoe*; Tce = *T. cristinae - T. monikensis*; Tps = *T. poppensis - T. douglasi*; Tcm = *T. californicum - T. shepardi*; Tpa = *T. podura - T. genevievae.*

| Sp | Tiss | FB_excl | total_excl | FB_kept | total_kept | FB_fisher_OR | FB_fisher_FDR |
|---|---|---|---|---|---|---|---|
| Tbi | WB | 8 | 758 | 582 | 19426 | 0.35 | **0.0021** |
| Tce | WB | 15 | 1069 | 1209 | 19757 | 0.22 | **0.0000** |
| Tcm | WB | 3 | 704 | 186 | 17661 | 0.40 | 0.1346 |
| Tpa | WB | 0 | 1383 | 73 | 22296 | 0.00 | **0.0366** |
| Tps | WB | 24 | 872 | 772 | 17862 | 0.63 | **0.0373** |
| Tbi | GN | 30 | 602 | 2182 | 16105 | 0.33 | **0.0000** |
| Tce | GN | 19 | 534 | 2282 | 16019 | 0.22 | **0.0000** |
| Tcm | GN | 14 | 500 | 1631 | 14412 | 0.23 | **0.0000** |
| Tpa | GN | 44 | 810 | 2765 | 16838 | 0.29 | **0.0000** |
| Tps | GN | 54 | 583 | 2106 | 15242 | 0.64 | **0.0029** |
| Tbi | LG | 22 | 654 | 289 | 15161 | 1.79 | **0.0256** |
| Tce | LG | 0 | 512 | 91 | 14761 | 0.00 | 0.0969 |
| Tcm | LG | 1 | 584 | 59 | 13733 | 0.40 | 0.5197 |
| Tpa | LG | 4 | 1602 | 95 | 15022 | 0.39 | 0.0809 |
| Tps | LG | 13 | 683 | 169 | 14497 | 1.64 | 0.1167 |

**Supplementary Table 10 | Number of male-biased genes excluded due to low expression in asexual females when using the asexual reference.** MB_excl = number of male-biased genes excluded, total_excl = total number of genes excluded, MB = number of male-biased genes in the main analysis, total = number of genes in the main analysis, fisher_OR = odd-ratio from a Fisher's exact test, fisher_FDR = FDR value from the Fisher's exact test. Tissue types are abbreviated as follows: WB = whole-body, RT = reproductive tract, and LG = legs. Species names are abbreviated by the sexual species name as follows (sexual species – asexual species): Tbi = *T. bartmani - T. tahoe*; Tce = *T. cristinae - T. monikensis*; Tps = *T. poppensis - T. douglasi*; Tcm = *T. californicum - T. shepardi*; Tpa = *T. podura - T. genevievae.*

| sp | Tiss | MB_excl | total_excl | MB_kept | total_kept | MB_fisher_OR | MB_fisher_FDR |
|---|---|---|---|---|---|---|---|
| Tbi | WB | 13 | 352 | 886 | 23653 | 0.99 | 1.0000 |
| Tce | WB | 37 | 618 | 1193 | 24292 | 1.23 | 0.3553 |
| Tcm | WB | 5 | 647 | 105 | 22607 | 1.67 | 0.3553 |
| Tpa | WB | 1 | 655 | 10 | 22943 | 3.51 | 0.3632 |
| Tps | WB | 9 | 531 | 493 | 21428 | 0.73 | 0.4936 |
| Tbi | GN | 147 | 604 | 3129 | 20237 | 1.76 | 0.0000 |
| Tce | GN | 86 | 514 | 2610 | 20487 | 1.38 | 0.0196 |
| Tcm | GN | 182 | 718 | 2300 | 19535 | 2.54 | 0.0000 |
| Tpa | GN | 165 | 752 | 2794 | 19073 | 1.64 | 0.0000 |
| Tps | GN | 144 | 515 | 3168 | 19169 | 1.96 | 0.0000 |
| Tbi | LG | 48 | 630 | 467 | 17876 | 3.07 | 0.0000 |
| Tce | LG | 7 | 373 | 118 | 17023 | 2.74 | 0.0341 |
| Tcm | LG | 14 | 629 | 138 | 16748 | 2.74 | 0.0032 |
| Tpa | LG | 10 | 835 | 145 | 16234 | 1.34 | 0.4009 |
| Tps | LG | 14 | 585 | 322 | 17455 | 1.30 | 0.4009 |

**Supplementary Table 11 | Number of female-biased genes excluded due to low expression in asexual females when using the asexual reference.** FB_excl = number of female-biased genes excluded, total_excl = total number of genes excluded, FB = number of female-biased genes in the main analysis, total = number of genes in the main analysis, fisher_OR = odd-ratio from a Fisher's exact test, fisher_FDR = FDR value from the Fisher's exact test. Tissue types are abbreviated as follows: WB = whole-body, RT = reproductive tract, and LG = legs. Species names are abbreviated by the sexual species name as follows (sexual species – asexual species): Tbi = *T. bartmani - T. tahoe*; Tce = *T. cristinae - T. monikensis*; Tps = *T. poppensis - T. douglasi*; Tcm = *T. californicum - T. shepardi*; Tpa = *T. podura - T. genevievae.*

| sp | Tiss | FB_excl | total_excl | FB_kept | total_kept | FB_fisher_OR | FB_fisher_FDR |
|---|---|---|---|---|---|---|---|
| Tbi | WB | 1 | 352 | 620 | 23653 | 0.11 | 0.0041 |
| Tce | WB | 3 | 618 | 1416 | 24292 | 0.08 | 0.0000 |
| Tcm | WB | 0 | 647 | 175 | 22607 | 0.00 | 0.0321 |
| Tpa | WB | 0 | 655 | 92 | 22943 | 0.00 | 0.2557 |
| Tps | WB | 11 | 531 | 798 | 21428 | 0.55 | 0.0697 |
| Tbi | GN | 21 | 604 | 2904 | 20237 | 0.22 | 0.0000 |
| Tce | GN | 13 | 514 | 2744 | 20487 | 0.17 | 0.0000 |
| Tcm | GN | 5 | 718 | 2019 | 19535 | 0.06 | 0.0000 |
| Tpa | GN | 38 | 752 | 3039 | 19073 | 0.28 | 0.0000 |
| Tps | GN | 35 | 515 | 2623 | 19169 | 0.46 | 0.0000 |
| Tbi | LG | 21 | 630 | 354 | 17876 | 1.71 | 0.0484 |
| Tce | LG | 0 | 373 | 87 | 17023 | 0.00 | 0.3349 |
| Tcm | LG | 0 | 629 | 46 | 16748 | 0.00 | 0.4786 |
| Tpa | LG | 3 | 835 | 99 | 16234 | 0.59 | 0.5257 |
| Tps | LG | 7 | 585 | 204 | 17455 | 1.02 | 0.8452 |

**Supplementary Table 12 | Numbers of sex-biased genes in each species when using the 10-species orthologs.** N FB, N MB, and N SB = number of female-, male-, or sex- biased genes respectively. SB dir = sex-bias direction, FDR = adjusted p-value for an excess of male- or female- biased genes. Species names are abbreviated as follows: Tbi = *T. bartmani*, Tce = *T. cristinae*, Tps = *T. poppensis*, Tcm = *T. californicum*, and Tpa = *T. podura*. Tissue types are abbreviated as follows: WB = whole-body, RT = reproductive tract, and LG = legs.

| **Species** | **Tissue** | **N FB** | **N MB** | **N SB** | **SB dir** | **FDR** |
|---|---|---|---|---|---|---|
| Tbi | WB | 124 (4.15) | 158 (5.28) | 282 (9.43) | MB | 0.0883 |
| | RT | 371 (13.20) | 520 (18.51) | 891 (31.71) | MB | **0.0000** |
| | LG | 73 (2.60) | 84 (2.99) | 157 (5.59) | MB | 0.4343 |
| Tce | WB | 188 (6.28) | 187 (6.24) | 375 (12.52) | FB | 0.9834 |
| | RT | 378 (13.27) | 429 (15.06) | 807 (28.34) | MB | 0.1263 |
| | LG | 26 (0.93) | 38 (1.35) | 64 (2.28) | MB | 0.1979 |
| Tps | WB | 174 (5.82) | 174 (5.82) | 348 (11.63) | NB | 1.0000 |
| | RT | 513 (18.16) | 558 (19.75) | 1071 (37.91) | MB | 0.2333 |
| | LG | 31 (1.10) | 49 (1.74) | 80 (2.85) | MB | 0.0883 |
| Tcm | WB | 57 (1.91) | 77 (2.58) | 134 (4.49) | MB | 0.1401 |
| | RT | 360 (12.75) | 424 (15.02) | 784 (27.77) | MB | 0.0636 |
| | LG | 18 (0.64) | 31 (1.11) | 49 (1.75) | MB | 0.1151 |
| Tpa | WB | 29 (0.97) | 14 (0.47) | 43 (1.44) | FB | 0.0636 |
| | RT | 545 (19.40) | 516 (18.36) | 1061 (37.76) | FB | 0.4343 |
| | LG | 23 (0.82) | 43 (1.54) | 66 (2.36) | MB | 0.0591 |

**Supplementary Table 13 |** Wilcoxon tests for expression shifts in sex-biased genes in asexual females when using 1-to-1 orthologs. ΔFB and ΔMB are the median shifts of female-biased and male-biased expression in asexual females respectively. Tte = *T. tahoe*, Tms = *T. monikensis*, Tdi = *T. douglasi*, Tsi = *T. shepardi*; Tge = *T. genevievae.* Tissue types are abbreviated as follows: WB = whole-body, RT = reproductive tract, and LG = legs.

| **Sp** | **Tissue** | **ΔFB** | **FB FDR** | **ΔMB** | **MB FDR** |
|---|---|---|---|---|---|
| Tte | WB | -0.217 | **$9.52 \times 10^{-05}$** | 0.035 | 0.5223 |
| Tms | WB | -0.224 | **$1.48 \times 10^{-11}$** | 0.214 | **$1.37 \times 10^{-13}$** |
| Tdi | WB | -0.627 | **$1.06 \times 10^{-36}$** | 0.318 | **$2.13 \times 10^{-14}$** |
| Tsi | WB | 0.175 | 0.2603 | -0.156 | 0.2425 |
| Tge | WB | -0.064 | 0.5583 | -0.049 | 0.5570 |
| Tte | RT | -0.171 | **$1.48 \times 10^{-11}$** | 0.161 | **$3.39 \times 10^{-12}$** |
| Tms | RT | -0.493 | **$1.22 \times 10^{-44}$** | 0.453 | **$1.12 \times 10^{-30}$** |
| Tdi | RT | -0.103 | **$3.30 \times 10^{-04}$** | 0.152 | **$2.03 \times 10^{-15}$** |
| Tsi | RT | 0.014 | 0.1189 | -0.023 | 0.0607 |
| Tge | RT | -0.191 | **$8.22 \times 10^{-12}$** | 0.145 | **0.0037** |
| Tte | LG | -0.507 | **$2.87 \times 10^{-05}$** | -0.134 | 0.4794 |
| Tms | LG | -0.244 | **0.0232** | 0.130 | **0.0484** |
| Tdi | LG | -0.827 | **$3.88 \times 10^{-04}$** | 0.506 | **$9.30 \times 10^{-06}$** |
| Tsi | LG | -0.073 | 0.4751 | -0.103 | 0.5408 |
| Tge | LG | -0.464 | **0.0069** | 0.446 | **0.0457** |

**Supplementary Table 14 |** FDR and gradients for the relationship between the number of species a gene is sex-biased in and the shift in expression in asexual females from a GLM (GLM_FDR) and a GLMM (GLMM_FDR) where gene ID was included as a random effect. Sex-bias = Female-biased (FB) or male-biased (MB) genes. Tissue types are abbreviated as follows: WB = whole-body, RT = reproductive tract, and LG = legs.

| Sex-bias | Tissue | Gradient | GLM_FDR | GLMM_FDR |
|---|---|---|---|---|
| FB | WB | 0.09099236 | 0.00252778 | 0.02017974 |
| FB | RT | 0.07658775 | 9.09E-09 | 3.53E-08 |
| FB | LG | 0.14406285 | 0.01665674 | 0.02218378 |
| MB | WB | -0.0942748 | 0.0012744 | 0.0012744 |
| MB | RT | -0.0440438 | 0.00223275 | 0.01255613 |
| MB | LG | -0.1891408 | 0.00631202 | 0.02218378 |

**Supplementary Table 15 | The effects of reproductive mode and sex-bias on dN/dS when modelled using a glmm approach.** LRT_bi = log-likelihood ratio test statistic for the binomial model, p_bi = p-value for the binomial model, LRT_nz = log-likelihood ratio test statistic for the gamma model, p_nz = p-value for the gamma model. Tissue types are abbreviated as follows: WB = whole-body, RT = reproductive tract, and LG = legs.

| Tissue | term | LRT_bi | p_bi | LRT_nz | p_nz |
|---|---|---|---|---|---|
| WB | rep mode | 268.221 | < 2e-16 | 62.858 | 2.22E-15 |
| WB | sex bias | 7.046 | 0.02951 | 6.549 | 0.03783 |
| WB | rep mode:sex bias | 0.69475 | 0.7065 | 4.8613 | 0.08798 |
| WB | OG_name | 436.736047 | < 2e-16 | 541.579247 | < 2e-16 |
| RT | rep mode | 266.388 | < 2e-16 | 75.116 | < 2e-16 |
| RT | sex bias | 6.787 | 0.03359 | 0.94 | 0.6251 |
| RT | rep mode:sex bias | 3.5051 | 0.1733 | 0.0082594 | 0.9959 |
| RT | OG_name | 424.217296 | < 2e-16 | 506.536595 | < 2e-16 |
| LG | rep mode | 261.564 | < 2e-16 | 71.029 | < 2e-16 |
| LG | sex bias | 11.477 | 0.003219 | 2.687 | 0.2609 |
| LG | rep mode:sex bias | 5.3507 | 0.06888 | 1.3921 | 0.4985 |
| LG | OG_name | 392.431522 | < 2e-16 | 498.694099 | < 2e-16 |

**Supplementary Table 16 | The effects of reproductive mode and sex-bias on dN/dS when modelled using a glmm approach, excluding female-biased genes.** LRT_bi = log-likelihood ratio test statistic for the binomial model, p_bi = p-value for the binomial model, LRT_nz = log-likelihood ratio test statistic for the gamma model, p_nz = p-value for the gamma model. Tissue types are abbreviated as follows: WB = whole-body, RT = reproductive tract, and LG = legs.

| Tissue | Term | LRT_bi | p_bi | LRT_nz | p_nz |
|---|---|---|---|---|---|
| WB | rep mode | 264.049 | < 2e-16 | 62.774 | 2.32E-15 |
| WB | sex bias | 1.103 | 0.2935 | 3.762 | 0.05242 |
| WB | rep mode:sex bias | 0.0180558 | 0.8931 | 3.1135 | 0.07765 |
| WB | OG_name | 405.501086 | < 2e-16 | 540.151184 | < 2e-16 |
| RT | rep mode | 242.251 | < 2e-16 | 62.571 | 2.57E-15 |
| RT | sex bias | 6.051 | 0.0139 | 0.107 | 0.7434 |
| RT | rep mode:sex bias | 1.4708 | 0.2252 | 0.019517 | 0.8889 |
| RT | OG_name | 333.263803 | < 2e-16 | 432.166011 | < 2e-16 |
| LG | rep mode | 264.966 | < 2e-16 | 71.214 | < 2e-16 |
| LG | sex bias | 9.469 | 0.00209 | 0.391 | 0.5317 |
| LG | rep mode:sex bias | 1.5662 | 0.2108 | 0.17648 | 0.6744 |
| LG | OG_name | 386.730801 | < 2e-16 | 491.822732 | < 2e-16 |

**Supplementary Table 17 | The effects of reproductive mode and sex-bias on dN/dS when modelled using a glmm approach, excluding male-biased genes.** LRT_bi = log-likelihood ratio test statistic for the binomial model, p_bi = p-value for the binomial model, LRT_nz = log-likelihood ratio test statistic for the gamma model, p_nz = p-value for the gamma model. Tissue types are abbreviated as follows: WB = whole-body, RT = reproductive tract, and LG = legs.

| Tissue | Term | LRT_bi | p_bi | LRT_nz | p_nz |
|---|---|---|---|---|---|
| WB | rep mode | 261.685 | < 2e-16 | 64.44 | 9.95E-16 |
| WB | sex bias | 6.315 | 0.01197 | 3.337 | 0.06772 |
| WB | rep mode:sex bias | 0.66007 | 0.4165 | 1.9975 | 0.1576 |
| WB | OG_name | 412.992912 | < 2e-16 | 520.666943 | < 2e-16 |
| RT | rep mode | 222.618 | < 2e-16 | 57.96 | 2.68E-14 |
| RT | sex bias | 0.382 | 0.54 | 0.982 | 0.3216 |
| RT | rep mode:sex bias | 2.6293 | 0.10 | 0.00258293 | 0.9595 |
| RT | OG_name | 291.599002 | < 2e-16 | 386.117859 | < 2e-16 |
| LG | rep mode | 252.98 | < 2e-16 | 67.225 | 2.42E-16 |
| LG | sex bias | 1.675 | 0.1956 | 2.32 | 0.1274 |
| LG | rep mode:sex bias | 3.7015 | 0.05436 | 1.214 | 0.2705 |
| LG | OG_name | 380.138803 | < 2e-16 | 489.202805 | < 2e-16 |

**Supplementary Table 18 |** Numbers of sex-biased genes in each species when using only an FDR threshold. N FB, N MB, and N SB = number of female-, male-, or sex- biased genes respectively. SB dir = sex-bias direction, FDR = adjusted p-value for an excess of male- or female- biased genes. Species names are abbreviated as follows: Tbi = *T. bartmani*, Tce = *T. cristinae*, Tps = *T. poppensis*, Tcm = *T. californicum*, and Tpa = *T. podura*. Tissue types are abbreviated as follows: WB = whole-body, RT = reproductive tract, and LG = legs.

| **Species** | **Tissue** | **N FB** | **MB_N** | **SB_N** | **SB dir** | **FDR** |
|---|---|---|---|---|---|---|
| Tbi | WB | 562 (3.68) | 643 (4.21) | 1205 (7.89) | MB | **0.0245** |
| | RT | 3830 (29.82) | 3735 (29.08) | 7565 (58.9) | FB | 0.2892 |
| | LG | 739 (6.12) | 842 (6.97) | 1581 (13.09) | MB | **0.0132** |
| Tce | WB | 1235 (8.32) | 1024 (6.9) | 2259 (15.22) | FB | **0.0000** |
| | RT | 1987 (16.23) | 1849 (15.1) | 3836 (31.33) | FB | **0.0314** |
| | LG | 83 (0.76) | 133 (1.21) | 216 (1.97) | MB | **0.0015** |
| Tps | WB | 697 (5.03) | 479 (3.46) | 1176 (8.49) | FB | **0.0000** |
| | RT | 3142 (26.75) | 3209 (27.32) | 6351 (54.06) | MB | 0.4108 |
| | LG | 159 (1.42) | 246 (2.19) | 405 (3.61) | MB | **0.0000** |
| Tcm | WB | 168 (1.2) | 121 (0.87) | 289 (2.07) | FB | **0.0088** |
| | RT | 1595 (14) | 1701 (14.93) | 3296 (28.94) | MB | **0.0701** |
| | LG | 50 (0.45) | 116 (1.04) | 166 (1.49) | MB | **0.0000** |
| Tpa | WB | 62 (0.45) | 5 (0.04) | 67 (0.48) | FB | **0.0000** |
| | RT | 2495 (22.54) | 2307 (20.84) | 4802 (43.38) | FB | **0.0098** |
| | LG | 76 (0.77) | 115 (1.16) | 191 (1.93) | MB | **0.0083** |

**Supplementary Table 19 |** Wilcox tests for expression shift in sex-biased genes in asexual females for when sex-biased genes are classed using only an FDR threshold. Tte = *T. tahoe*, Tms = *T. monikensis*, Tdi = *T. douglasi*, Tsi = *T. shepardi*; Tge = *T. genevievae.* Tissue types are abbreviated as follows: WB = whole-body, RT = reproductive tract, and LG = legs.

| **Sp** | **Tissue** | **ΔFB** | **FB FDR** | **ΔMB** | **MB FDR** |
|---|---|---|---|---|---|
| Tte | WB | -0.255 | **$7.29 \times 10^{-28}$** | 0.136 | **$1.80 \times 10^{-05}$** |
| Tms | WB | -0.164 | **$1.33 \times 10^{-38}$** | 0.211 | **$6.82 \times 10^{-34}$** |
| Tdi | WB | -0.848 | **$3.50 \times 10^{-193}$** | 0.749 | **$3.73 \times 10^{-65}$** |
| Tsi | WB | -0.283 | **$9.69 \times 10^{-06}$** | 0.223 | **$3.04 \times 10^{-06}$** |
| Tge | WB | -0.184 | 0.0755 | -0.252 | 0.2587 |
| Tte | RT | -0.126 | **$3.73 \times 10^{-51}$** | 0.162 | **$4.26 \times 10^{-48}$** |
| Tms | RT | -0.368 | **$4.92 \times 10^{-113}$** | 0.351 | **$3.56 \times 10^{-134}$** |
| Tdi | RT | -0.063 | **$1.07 \times 10^{-17}$** | 0.087 | **$1.55 \times 10^{-11}$** |
| Tsi | RT | 0.009 | **$3.76 \times 10^{-04}$** | -0.008 | 0.0541 |
| Tge | RT | -0.199 | **$2.16 \times 10^{-61}$** | 0.087 | 0.4055 |
| Tte | LG | -0.252 | **$1.55 \times 10^{-37}$** | -0.087 | **$2.03 \times 10^{-06}$** |
| Tms | LG | -0.175 | **0.0024** | 0.051 | 0.1464 |
| Tdi | LG | -0.590 | **$1.44 \times 10^{-17}$** | 0.314 | **$1.41 \times 10^{-12}$** |
| Tsi | LG | -0.291 | **$6.68 \times 10^{-05}$** | -0.070 | 0.7539 |
| Tge | LG | -0.503 | **$4.15 \times 10^{-07}$** | 0.274 | **$9.61 \times 10^{-04}$** |
